# Cytokine and phenotypic cell profiles in human cutaneous leishmaniasis caused by *Leishmania donovani*

**DOI:** 10.1101/2022.06.16.496495

**Authors:** Hiruni Wijesooriya, Nilakshi Samaranayake, Nadira D. Karunaweera

## Abstract

**Background:** The innate immune mediators are likely to influence the clinical phenotype of leishmaniasis by primary responses which limit or facilitate the spread of the parasite, as well as by modulating adaptive immunity. This study investigated the response of key innate immune cells in a focus which regularly reports localised cutaneous leishmaniasis (LCL) caused by *Leishmania donovani*, a species which typically causes visceral disease.

**Methods:** Peripheral blood mononuclear cell (PBMC) derived macrophage and dendritic cell responses to soluble *Leishmania* antigen (SLA) were compared between patients with LCL and healthy controls from endemic and non-endemic areas. Inflammatory mediators produced by macrophages (TNF-α, NO, TGF-β and IL-10) and dendritic cells (IL-12p70, IL-10) and cell surface markers of macrophage polarization, activation and maturation were determined at 24h, 48h and 72h by Enzyme-linked immunosorbent assay (ELISA) and flowcytometry.

**Results:** Patient derived macrophages and dendritic cells produced higher levels of both pro and anti-inflammatory mediators compared to controls (p<0.05) with the best discrimination for active disease observed at 72h. Data demonstrated an early activation of macrophages and a subsequent pro-inflammatory bias, as indicated by temporal profiles of TNF-α/TGF-β and TNF-α/IL-10 ratios and higher proportions of classical (M1) macrophages. Higher TGF-β levels were observed in cells from patients with ulcerated or persistent lesions. Immune responses by cells derived from controls in endemic and non-endemic regions did not differ significantly from each other.

**Conclusions:** The overall immunophenotypic profile suggests that LCL observed in the country is the result of a balancing immune response between pro-inflammatory and regulatory mediators. The mediators which showed distinct profiles in patients warrant further investigation as potential candidates for immunotherapeutic approaches. A comparison with visceral leishmaniasis caused by the same species, would provide further evidence on the differential role of these mediators in the resulting clinical phenotype.

## Introduction

Leishmaniasis is a vector-borne parasitic disease, caused by protozoan parasites of the genus *Leishmania*. The clinical picture of leishmaniasis is heterogeneous with a wide spectrum of human diseases, including cutaneous (CL), mucocutaneous and visceral (VL) forms. It is endemic in 98 countries with an estimated annual incidence of 30 000 cases of VL and more than 1 million cases of CL [1]. The cutaneous form of leishmaniasis itself has diverse presentations; the most common being localized cutaneous leishmaniasis (LCL). Further, in areas endemic for *Leishmania* transmission, approximately 10% of individuals may have evidence of exposure to parasite but lack disease pathology [2] and are classified as those with subclinical infections.

The outcome of infection in leishmaniasis is complex, depending not only on the parasite species but also on the immune status of the host. The clear dichotomy of Th1/Th2 reactions seen in experimental models, with a Th1 type protective response being associated with localised cutaneous disease, is not always observed in leishmaniasis in human hosts. Evidence from more recent studies suggests that the innate immune response plays a pivotal role in the outcome in *Leishmania* infections. This response would act both in controlling parasite growth during the early stages of infection as well as in driving the cytokine microenvironment in which parasite-specific T cells are primed [3-6].

Macrophages and dendritic cells; two main cell types of the innate immune system, play a decisive role in the initial interactions between the parasites and the immune system of the host. Macrophages serve as host cells as well as the effector cells which finally eliminate the parasite, whereas both are accessory cells that present parasite antigens, deliver co-stimulatory signals and secrete cytokines modulating the subsequent immune response. It is noteworthy that macrophages generally lack the ability to induce the primary stimulation of specific T cells, which is carried out by dendritic cells, thus linking the innate and adaptive immune systems.

Sri Lanka, an island nation in South Asia, mostly reports LCL with a few reports of mucosal [7] and visceral [8, 9] disease. The causative agent of all clinical disease types in Sri Lanka has been identified as *Leishmania donovani* MON 37 [10, 11]. *L. donovani* is usually associated with VL in the Old World and this species has been found to cause CL in only a few other foci; in Kenya [12], Yemen [13] and in the Himalayan region of northern India [14]. The pathogenic mechanisms which limit this usually visceralizing species to localised cutaneous lesions remain largely unknown, with only limited reports from Sri Lanka on possible contributory host factors based on experimental models [15] and human studies [16-19].

The objective of this study was to characterize the innate cellular immune responses associated with locally acquired cutaneous leishmaniasis due to *L. donovani*. We hypothesized that distinct profiles of the early immune response determined the clinical phenotype of localized cutaneous leishmaniasis observed in Sri Lanka. We aimed to investigate these immunological responses in LCL in relation to patients, non-endemic controls as well as endemic controls, with the latter likely to have been exposed but not developed symptomatic infections.

## Materials and Methods

### Ethics approval and consent to participate

The study was conducted in compliance with the principles of Declaration of Helsinki [20]. All participants provided informed written consent and study protocols were approved by the Ethics Review Committee of Faculty of Medicine, University of Colombo, Sri Lanka.

### Study locations, participants and sample collection

Altogether sixty patients with locally-acquired cutaneous leishmaniasis were recruited for the study from the leishmaniasis clinic held weekly at the Department of Parasitology, Faculty of Medicine, University of Colombo and dermatology clinics in selected Base Hospitals in endemic areas. The exclusion criteria consisted a history of inflammatory / immunosuppressive medical conditions. Details of the patients were collected using a pretested questionnaire by directly interviewing and examining the patients. Diagnosis was confirmed by direct microscopy and/ or culture of lesion material obtained by aspirates or slit skin smears. 5ml of peripheral venous blood was collected by venipuncture to EDTA tubes from each participant. All patients were routinely tested by rK39 immunochromatographic rapid diagnostic test to exclude serological evidence of visceral involvement.

Sixty endemic controls and thirty non-endemic controls, within the same age range and with the same gender distribution as the patients, with no history of inflammatory /immune suppressive medical conditions comprised the comparison group. Healthy non endemic controls were residents from areas which are not endemic for leishmaniasis in Sri Lanka. Healthy household contacts of diagnosed patients with no past history suggestive of leishmaniasis were recruited as endemic controls (Fig. 1).

**Fig 1.**
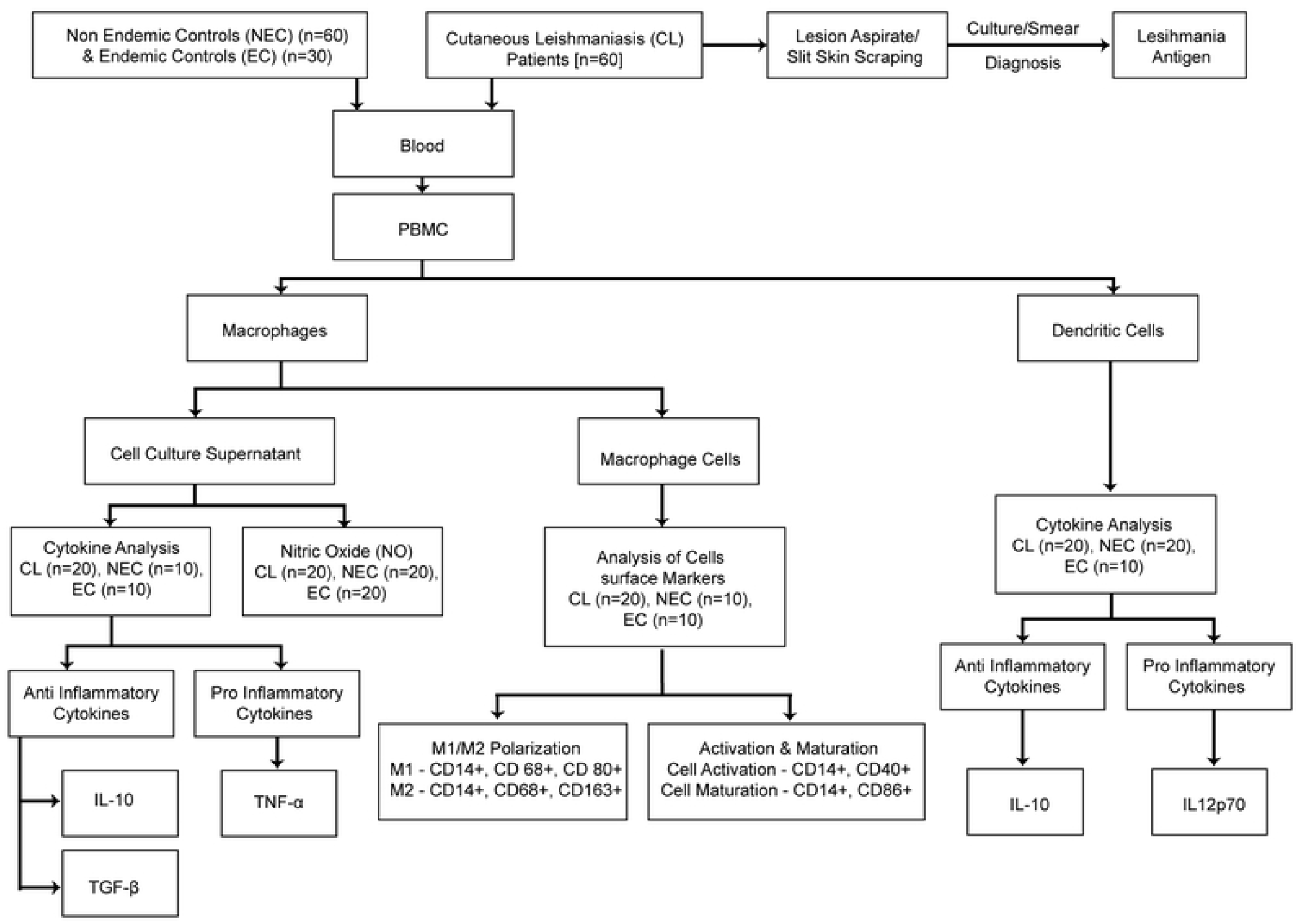
Participant recruitment and sample analysis plan.

### Preparation of soluble *Leishmania* antigen (SLA)

*L. donovani* parasites were cultured in complete M199 (Gibco, USA) media (Hank’s salts, L-Glutamine, 25mM HEPES, L-Amino Acids) with5% heat inactivated fetal bovine serum (Sigma, USA) and 0.1% gentamycin. Antigen extracts were prepared from stationary phase promastigotes by 6 cycles of freezing and thawing as described elsewhere [21]. The antigen concentration was measured by Lowry’s protein quantification method [22] and stored at -70°C until use.

### Generation of human monocyte-derived macrophages and dendritic cells *in-vitro*

Peripheral Blood Mononuclear Cells (PBMC) were isolated from EDTA blood by density gradient centrifugation using 30% of Ficoll-Paque (GE Life Sciences, country). Cells were incubated at 37°C in 5% CO_2_ and monocytes were separated by adherence after 24 hours and cultured in complete RPMI1640(Gibco, USA) supplemented with 5% human heat-inactivated serum (Sigma, USA),10% of fetal calf serum (FCS)(Sigma, USA) and100 U/mL Penicillin-Streptomycin(Sigma, USA)in 24 well plates (1 × 106cells/well) [23]. The adherent cells displayed characteristics of monocyte derived macrophages after 5-6 days.

Dendritic cells were generated by culturing the monocytes separated by adherence at 24hrs in complete RPMI supplemented with 200ng/ml recombinant human granulocyte macrophage colony stimulating factor (GMCSF) and 100ng/ml recombinant IL-4 for 6 days [24].

### Macrophage and dendritic cell stimulation with SLA

PBMC derived macrophages/dendritic cells were stimulated with SLA at a concentration of 50ug/ml for 6 days in culture. Uninfected cells and Con –A (50Ug/ml) were used as negative and positive controls respectively. Macrophage cell culture supernatants were harvested at 24h, 48h and 72h time intervals and stored at -20°C for analysis of cytokines and NO levels. Dendritic cells were detached using ice cold PBS/EDTA solution at 24h, 48h and 72h time intervals and used for flowcytometric analysis.

### Cytokine and nitric oxide assays

Supernatants of macrophage cultures from LCL patients, endemic and non-endemic controls were assayed for TNF-α, TGF-β & IL-10 after 24, 48 and 72 hours of stimulation by ELISA according to manufacturer’s instructions(R&D Systems, USA). The sensitivity of the cytokine assays were 1.6pg/ml(TNF-α), 4.61pg/ml (TGF-β) and 3.9 pg/ml (IL-10). Nitrite (NO_2−_)accumulation in the cell culture supernatants from all the subjects were measured as an indicator of NO production using a standard Griess reagent kit (Molecular Probes, USA; minimum detection level 1.0 μM).

Intracellular cytokine staining (ICS) was used to assess cytokine production in dendritic cells. Fixation and permeabilization of the cells were conducted with fixation buffer (BioLegend, USA) and 10% permeabilization wash buffer (BioLegend, USA). FITC-conjugated anti-CD11c, APC-conjugated anti-IL-12p70, and PE-conjugated anti-IL-10 were used for the cytokine detection and finally the cells were dislodged in PBS containing 2% formaldehyde. All flowcytometric data was acquired with Cube 8 (Partech) flow cytometer and analyzed using FCS Express 4 (De Novo) software.

### Flowcytometric analysis of cell surface markers

Expression of selected cell surface receptors on macrophages was assayed by flowcytometry. After washing with PBS, cells were incubated at 4°C with either CD14-PE (phycoerythrin), CD68-FITC (fluorescein), CD80-APC (allophycocyanin) and CD163-Per-CP (peridinin chlorophyll) or CD 14-PE, CD40-FITC and CD86-Per-CP for staining, washed again and acquired and analyzed as above. Single stained cells and unstained cells were used to confirm the specificity of the binding of the antibody of interest and to rule out non-specific receptor binding. All antibodies were purchased from BioLegend, USA.

### Statistical analysis

Levels of inflammatory biomarkers are presented as medians and interquartile ranges (IQRs). Differences in these biomarker levels between the patients and the control groups were assessed using non-parametric Mann–Whitney U test or Kruskal-Wallis test followed by Dunn’s multiple comparison test, as appropriate. Receiver Operating Characteristic (ROC) curve analysis was conducted and area under the curve (AUC) was compared to determine the ability of the inflammatory markers, individually and in combination, to differentiate patients from controls. Analysis was performed using IBM SPSS version 20 and GraphPad Prism version 7 (GraphPad Software, Inc., La Jolla, CA, USA). All statistical tests were two-tailed and a probability (*P*) value of less than 0.05 was considered statistically significant.

## Results

### Characteristics of the study population

The study population altogether consisted of 60 patients and 90 controls. The socio demographic and clinical characteristics of the patients are summarized in Table 1. Patients consisted of 40 (66.7) males and 20 (33.3) females with a median age was 47 years (range 19–60 years). A majority of the lesions were either papules or nodules (37/60; 61%) with other lesion types seen in fewer numbers (Table 1). None of the patients had similar lesions in the past. All except nine patients had single lesions (51/60). None of the patients showed evidence of visceralization.

**Table 1.**
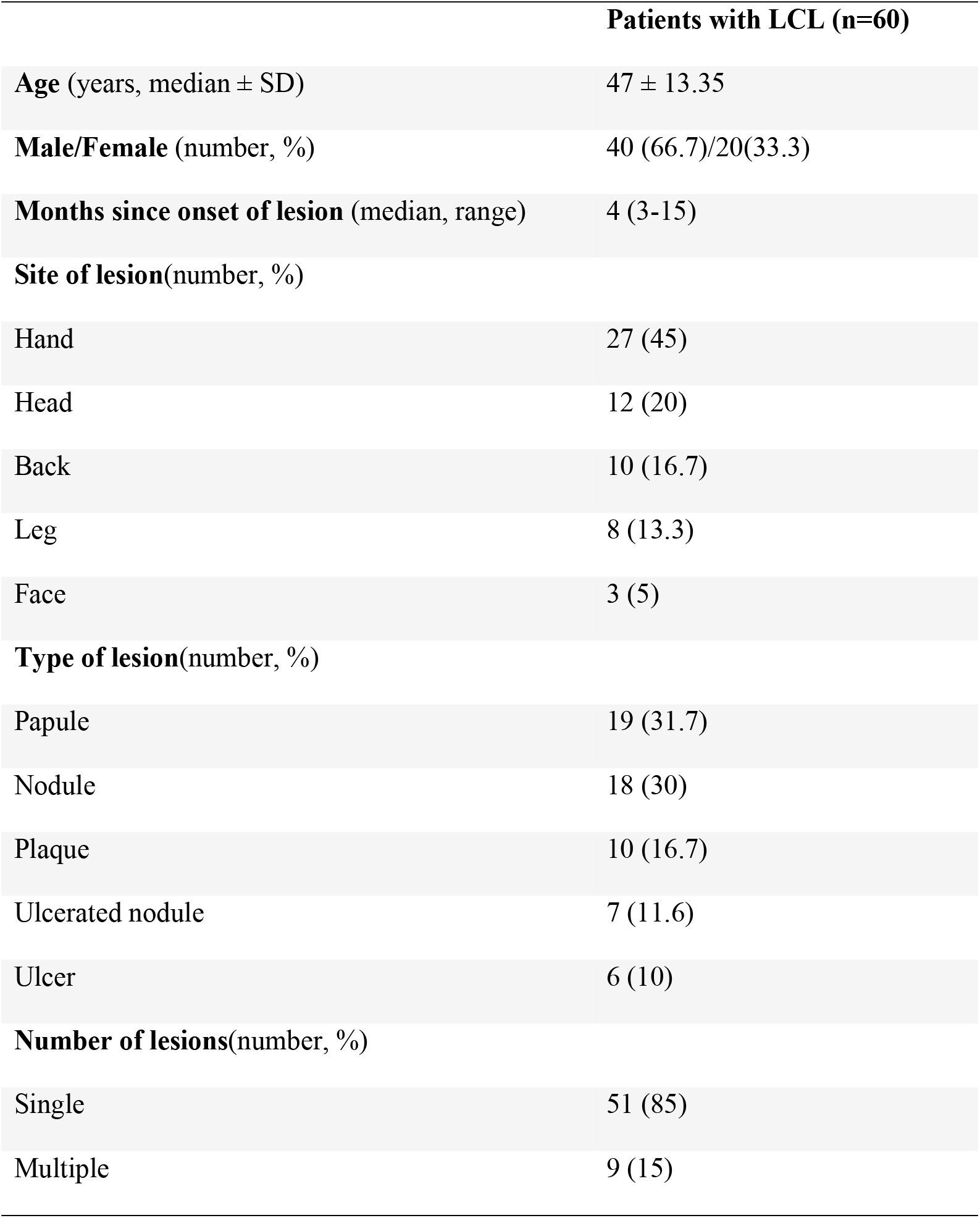
Sociodemographic and clinical characteristics of patients enrolled in the study.

### Patient derived macrophages produced a mixed inflammatory pattern after *in vitro* stimulation with SLA

We determined the temporal immune response of patient and control derived macrophages through the comparison of the levels of selected cytokines and NO in cell culture supernatants at 24, 48 and 72 hours (Fig 2).We observed a relative increase in production of TNF-α by patient derived cells which was significantly higher at 48 and 72 hours (when compared with both endemic and non-endemic controls; p<0.01).

**Fig 2.**
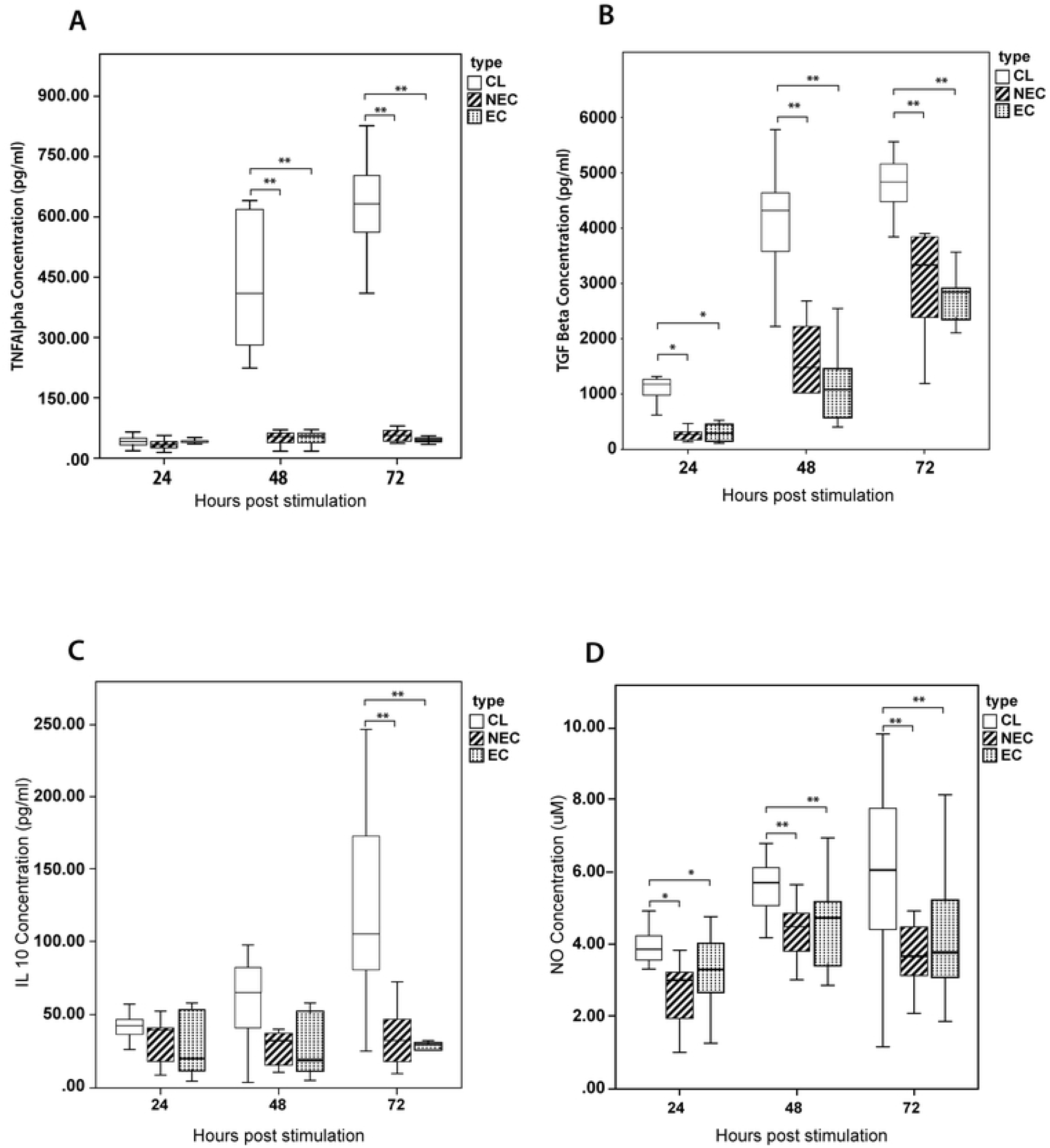
Comparison of selected inflammatory markers secreted by PBMC derived macrophages stimulated with SLA. Monocyte derived-macrophages from patients with LCL (n=20), healthy non endemic controls (n=10) and healthy endemic controls (n=20) were stimulated with *L. donovani* antigen (50µg/ml). The supernatant was harvested at 24, 48 & 72 hours and (A) TNF-α (B) TGF-β (C) IL-10 and (D) NO levels were determined by ELISA and Griess test respectively (in duplicate). The box represents the IQR (i.e. the middle 50% of the observations) with the horizontal line representing the median. The whiskers represent the main body of the data, indicating the range of the data. For statistical analysis, nonparametric Kruskal-Wallis test followed by Dunn’s multiple comparison test was used (*p<0.05; **p<0.01; ***p<0.001).

Whereas IL-10 secretion by patient derived cells showed a marked increase at 72hrs when compared to both endemic and non-endemic controls (p<0.01), both IL-10 and TNF-α levels in control groups remained relatively low throughout.

Secretion of TGF-β by patient derived cells was higher compared to control derived cells at all time points with a sharp rise in levels observed at 48hrs which persisted at 72hrs. The differences in cytokine levels were significant at all 24h (vs endemic controls, p=0.027; vs non-endemic controls, p= 0.003), 48h (vs endemic controls, p<0.01; vs non-endemic controls, p= 0.001) and 72h (vs endemic controls, p=0.008; vs non-endemic controls, p<0.01) time points.

Nitric oxide (NO) levels showed a pattern of differences similar to TGF-β with secretion by patient derived cells being higher at 24h (vs endemic controls, p=0.03; vs non-endemic controls, p=0.009), 48h (vs endemic controls, p=0.002; vs non-endemic controls, p<0.01) and 72h time points (vs endemic controls, p=0.038; vs non-endemic controls, p=0.009).

None of the cytokine combinations showed a strong correlation of secretion levels at any of the three time points (25).

### Patient derived dendritic cells produced a mixed inflammatory pattern after *in vitro* stimulation with SLA

A similar temporal analysis was carried out on the immune response by dendritic cells derived from patients and controls after in vitro stimulation with SLA (Fig 3A&3B). IL-10 and IL-12p70 intra cellular markers were used in combination with CD11c to determine the intracellular cytokine production by dendritic cells.

**Fig 3.**
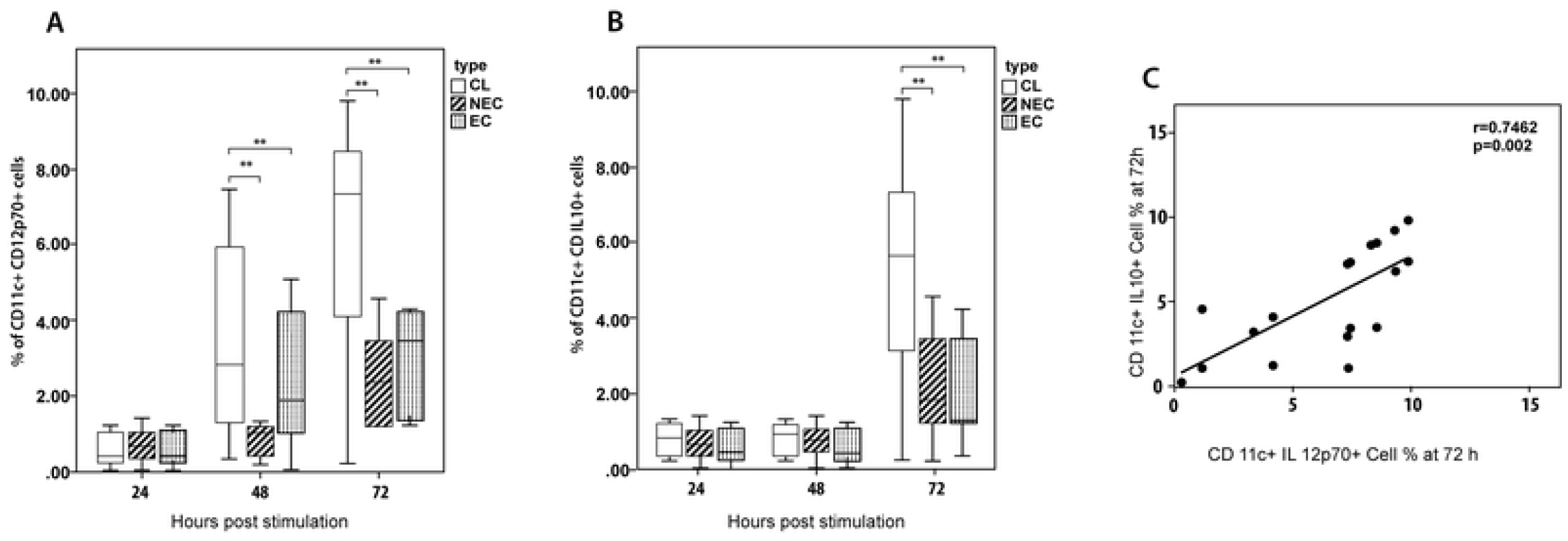
Comparison of selected cytokines production by PBMC derived dendritic cells stimulated with SLA. Dendritic cells from LCL (n=20), healthy non endemic controls (n=10) and healthy endemic controls (n=20) were stimulated with *L* .*donovani* antigen (50µg/ml). The supernatant was harvested at 24, 48 and 72 hours and intracellular (A) IL-12p70 and (B) IL-10 levels were determined by flowcytometry(in duplicate). The box represents the IQR (i.e. the middle 50% of the observations) with the horizontal line representing the median. The whiskers represent the main body of the data, indicating the range of the data. Cytokine levels were compared using nonparametric Kruskal-Wallis test followed by Dunn’s multiple comparison test (*p<0.05; **p<0.01; ***p<0.001). (C) IL-12p70 and IL-10 levels showed a strong positive correlation at 72 h (Spearman correlation test, p< 0.05).

The percentage of CD11c+IL-12p70+ in patient derived cells showed a steady increase over time unlike the cells derived from controls and this increase was significant at both 48h (vs endemic controls, p=0.026; vs non-endemic controls, p=0.027) and 72h (vs endemic controls, p=0.042; vs non-endemic controls, p=0.01) time points. The percentage of CD11c+IL10+ cells showed a marked rise in patient derived cells at 72 hours and was significantly higher compared to cells derived from both endemic (p=0.018) and non-endemic(p=0.027) controls.

IL-12p70 and IL-10 levels showed a strong positive correlation at 72 h (Fig 3C) (r=0.7462, p=0.0002).

### Pro inflammatory vs. regulatory cytokines and utility as diagnostic biomarkers

The temporal dynamics of inflammatory mediators produced by patient derived cells showed a distinct pattern of early responses to parasite antigen (Fig 4 A-F). We further analysed the cytokine responses at 72hrs, the time point at which most of the significant differences were observed. Macrophage derived pro-inflammatory to anti-inflammatory cytokine ratios were seen to be significantly higher in patients compared to controls (TNF-α to IL-10, p=0.008; TNF-α to TGF-β, p< 0.001)(Fig 4 G-H).While the proportion of dendritic cells producing IL-12p70 and IL-10 also favoured a pro inflammatory response as indicated by IL-12p70 /IL-10 ratio, the differences between the patients and the controls was not significant.

**Fig 4.**
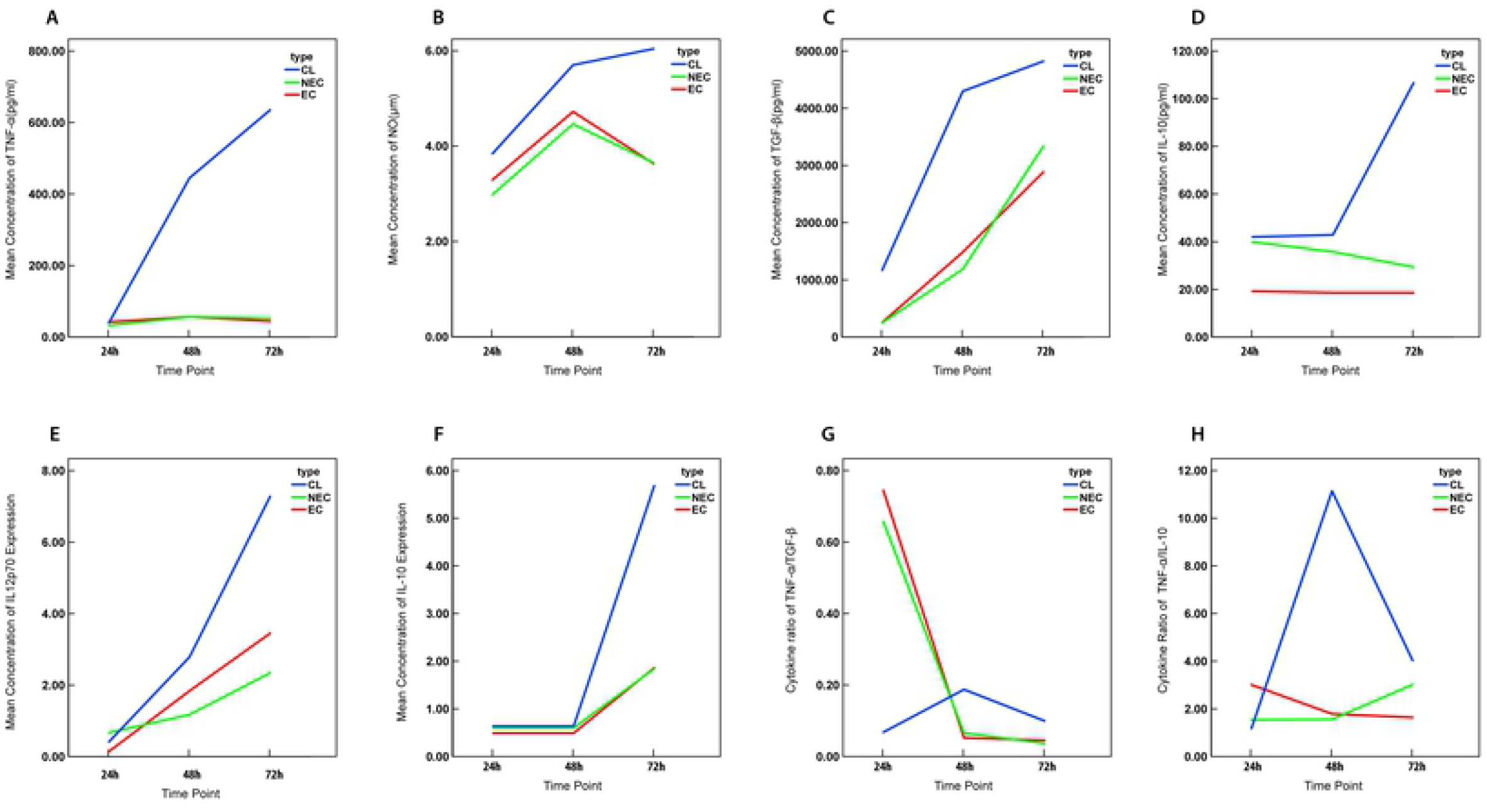
Change in mean concentration of inflammatory markers and their ratios over time. Line graphs representing the mean production of (A)TNF-α (B)NO (C)TGF-β (D) IL-10 in macrophages and (E)IL-12p70 (F)IL-10 in dendritic cells stimulated with *L. donovani* antigen (50µg/ml) and the ratio of the mean concentration of (G)TNF-α /TGF-β (H)TNF-α/ IL-10 at 24, 48 and 72 hour time intervals.

We performed receiver operating characteristic (ROC) curve analysis in order to assess the predictive value of the levels of the inflammatory markers to accurately identify patients with active disease. In line with the profile of significant differences between patients and controls, the ROC curves showed best discrimination at 72 hours for all 3 cytokines (TNF-α, IL-10 and TGF-β) produced by macrophages(Table 2)(Fig 5).

**Table 2.**
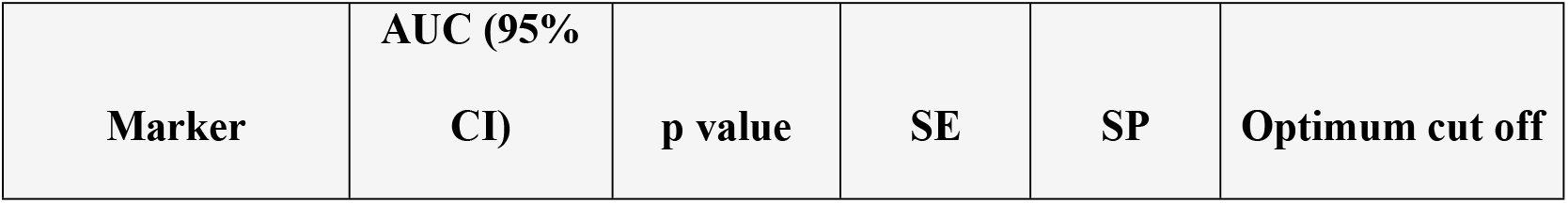
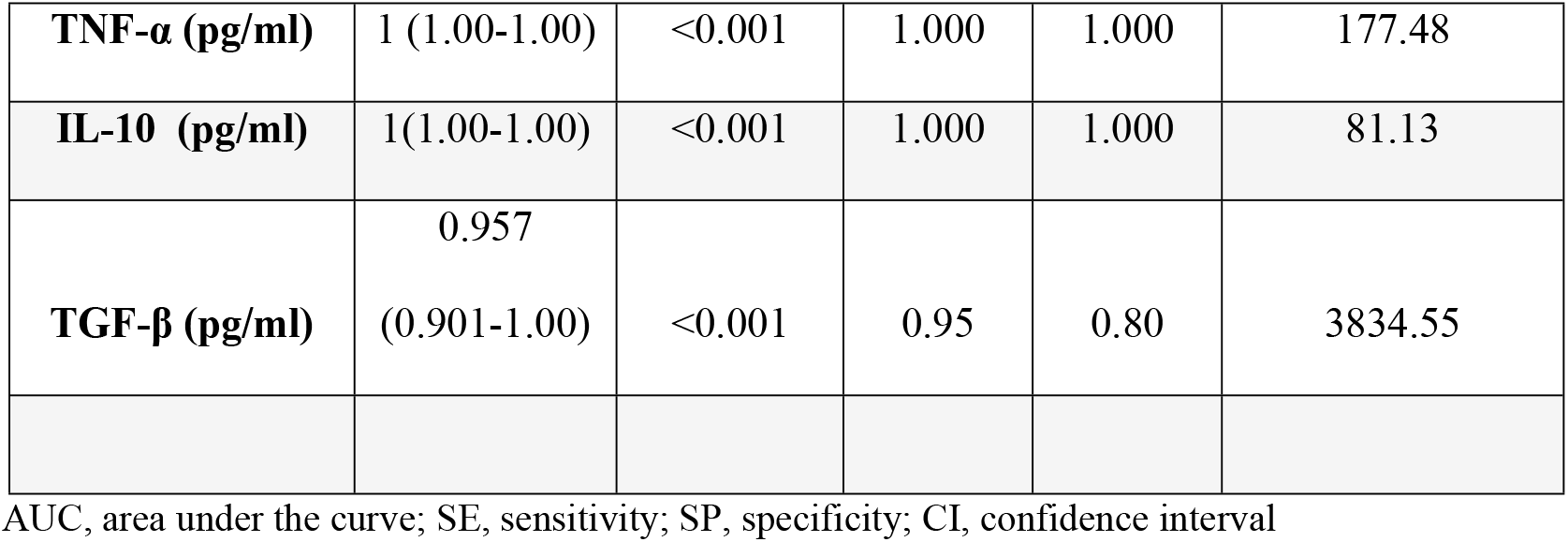
Performance of tested analytes as biomarkers to identify active disease status.

| Marker | AUC (95% CI) | p value | SE | SP | Optimum cut off |
| --- | --- | --- | --- | --- | --- |
| **TNF-α (pg/ml)** | 1 (1.00-1.00) | <0.001 | 1.000 | 1.000 | 177.48 |
| **IL-10 (pg/ml)** | 1(1.00-1.00) | <0.001 | 1.000 | 1.000 | 81.13 |
| **TGF-β (pg/ml)** | 0.957<br>(0.901-1.00) | <0.001 | 0.95 | 0.80 | 3834.55 |

AUC, area under the curve; SE, sensitivity; SP, specificity; CI, confidence interval

**Fig 5.**
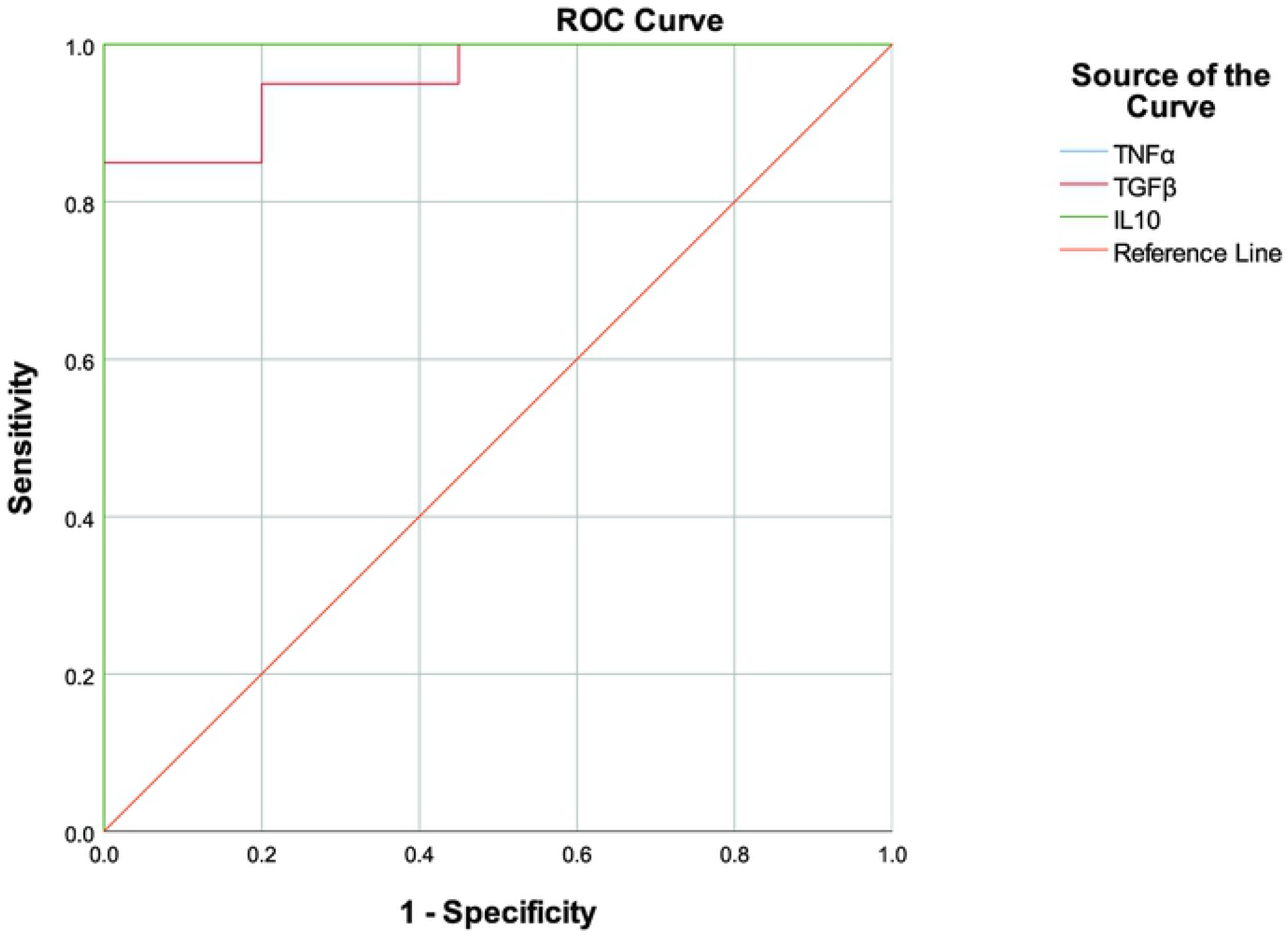
ROC curves for inflammatory markers produced by macrophages. ROC curves were calculated for TNF-α, NO, TGF-β and IL-10 produced by macrophages which showed significant differences in mean concentration levels in the culture supernatant. Detailed information on the AUCs is shown in Table 2. Higher values on the y-axis correspond to higher sensitivity, whereas lower values on the x-axis correspond to higher specificity.

The endemic and non-endemic controls were pooled for the above analyses since significant differences were not observed in cytokine levels between these two groups of controls.

### Patient derived macrophages displayed M1 polarization after stimulation with SLA

Macrophage polarization into M1or M2 subsets was identified by expression of CD14, CD68, CD80 and CD163 markers. CD14 and CD68 markers are expressed by monocytes and macrophages and M1 and M2 cells are known to predominantly express CD80 and CD163 respectively. The frequency of CD14+CD68+CD80+ macrophages were significantly higher in LCL patients when compared to healthy individuals at 48h (vs endemic controls, p=0.012; vs non-endemic controls, p=0.013) and 72h (vs endemic controls, p=0.045; vs non-endemic controls, p=0.006) time points (Figs 6B and 6C). But in contrast the fraction of CD14+CD68+ CD163+ cells were low in the same subjects at all the time points (26) suggesting a M1 polarization, which is associated with pro-inflammatory microbicidal responses.

**Fig 6.**
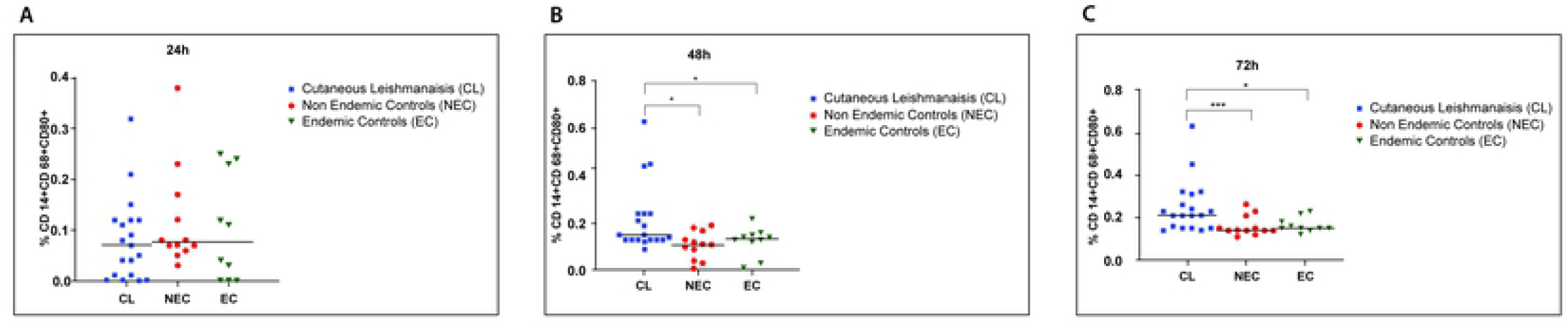
Distribution of M1 macrophages in patients and controls following stimulation with SLA. Monocyte derived-macrophages from LCL (n=20), healthy non endemic controls (n=10) and healthy endemic controls (n=20) were stimulated with *L*.*donovani* antigen (50µg/ml). Expression of CD14, CD68 and CD80 were analysed by flowcytometry at A) 24h, B) 48h and C) 72h to determine M1 polarization (in duplicate). In the vertical scatter plot each symbol represents a different subject. Horizontal bars represent the median.For statistical analysis, nonparametric Kruskal-Wallis test followed the Dunn’s multiple comparison test was used was used (*p<0.05; **p<0.01; ***p<0.001).

### *Leishmania donovani* antigen promotes early activation but not maturation of patient derived macrophages

CD40+ and CD86+ cells were gated on CD14+ cells in order to determine the cell activation and maturation respectively of macrophages upon stimulation with SLA. CD14+CD40+ cell population was significantly higher at the 24h time point in LCL patients than the controls (vs endemic controls, p=0.006;vsnon-endemic controls, p=0.011) suggesting early activation of the monocyte/macrophage system in this group (Fig 7A) with comparable levels of activation of patient and control derived cells at 48 and 72 hours (Fig 7B and C). However, the CD14+CD86+ cell population did not differ between the patients and the controls throughout the time course (27).

**Fig 7.**
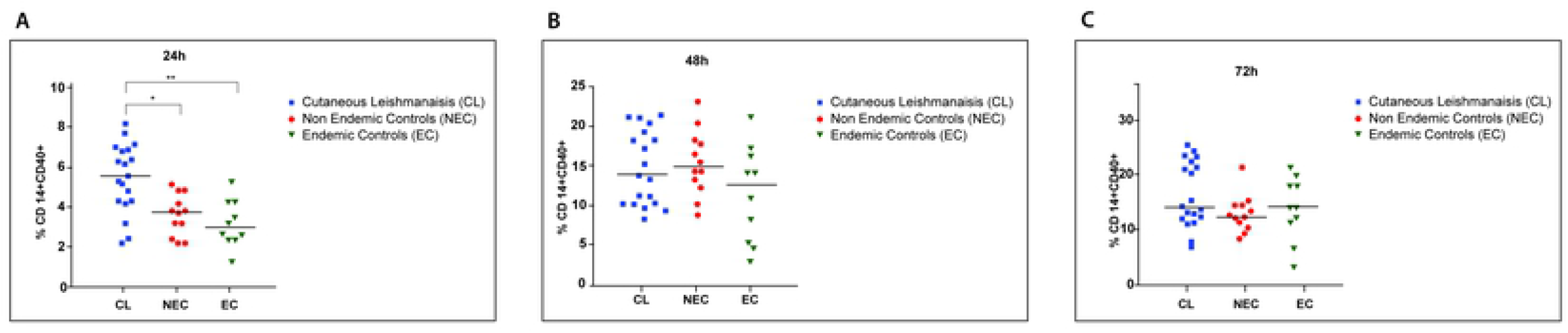
Distribution of activated macrophages over time in patients and controls following stimulation with SLA. Monocyte derived macrophages from LCL (n=20), healthy non endemic controls (n=10) and healthy endemic controls (n=20) were stimulated with *L*.*donovani* antigen (50µg/ml). Percentage of activated cells was determined at A) 24h, B) 48h and C) 72h by analyzing expression of CD14 and CD40 by flowcytometry. In the vertical scatter plot each symbol represents a different subject. Horizontal bars represent the median.For statistical analysis, nonparametric Kruskal-Wallis test followed the Dunn’s multiple comparison test was used was used (*p<0.05; **p<0.01; ***p<0.001).

### TGF-β secretion by macrophages is associated with severity of the cutaneous lesions

The immune response was evaluated in relation to lesion characteristics with late stage (lesion present for more than six months) and/or ulcerated lesions considered as features of severity. Cells from patients with both acute and chronic lesions consistently showed a sharp increase in production of TGF-β at 48 and 72hrs, with those from late stage lesions showing significantly higher levels at 48hrs (p=0.041)(Fig 8A).

**Fig 8.**
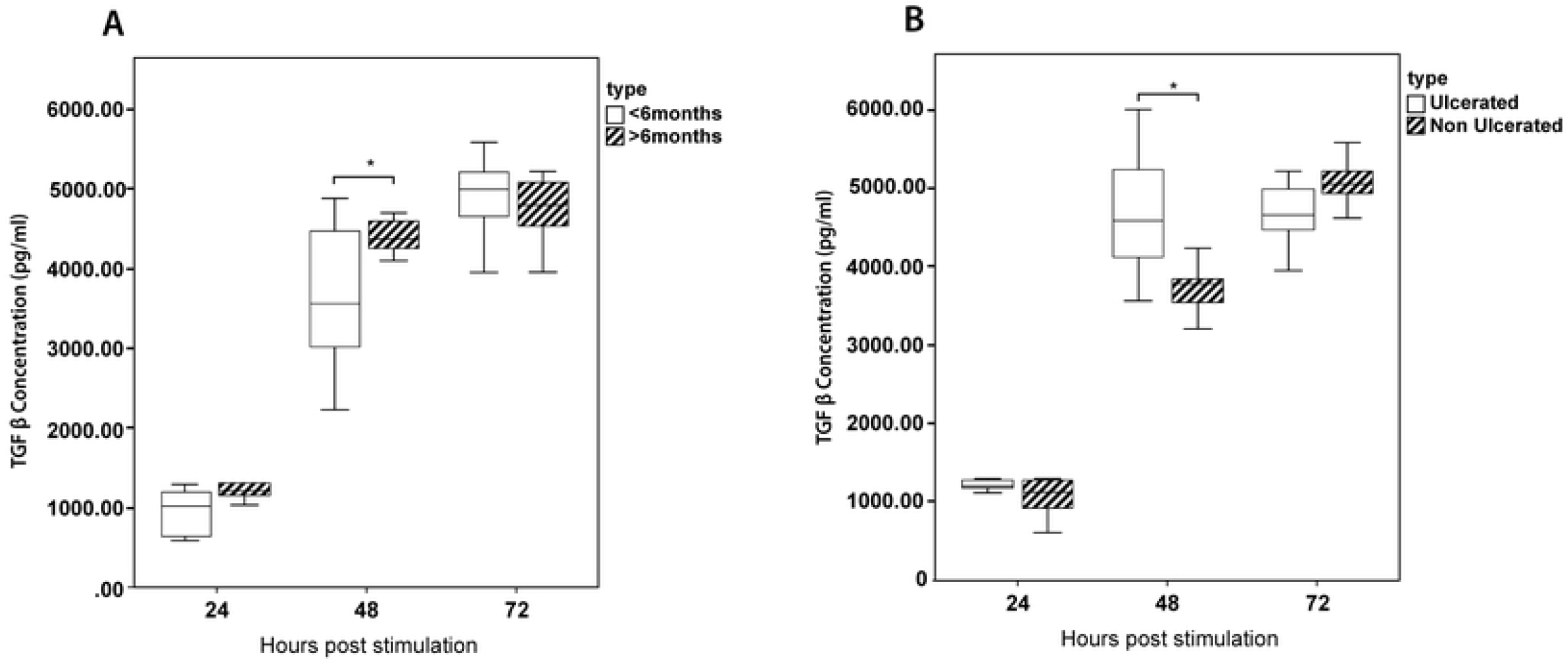
Kinetics of TGF-β in relation to lesion type and duration. The production of TGF-β was compared between patients with (A) acute (lesions less than six months/chronic lesions (lesions more than 6 months) and (B) ulcerated/non ulcerated lesions. The box represents the IQR (i.e. the middle 50% of the observations) with the horizontal line representing the median.The whiskers represent the main body of the data, indicating the range of the data. For statistical analysis, nonparametric Kruskal-Wallis test followed by the Dunn’s multiple comparison test was used (*p<0.05; **p<0.01; ***p<0.001).

Comparative evaluation with lesion type showed a similar trend with cells from patients with ulcerated lesions showing significantly higher levels of TGF-β at 48hrs (p=0.009) (Fig 8B). IL-10, TNF-α and NO levels did not differ significantly with the duration of the lesion or presence of ulceration. A similar analysis of IL-10 and IL12p70 secreted by dendritic cells also did not show any significant differences (28).

The same pattern of MI polarization was retained in acute lesions (Fig S1A). While cells from patients with chronic lesions showed a marked increase in M2 macrophages at the same time point when compared to acute lesions, this difference was not statistically significant (p>0.05) (Fig S1B). The presence of activated macrophages also appeared to favour ulcerated lesions (Fig S2).

## Discussion

A role for cytokines and phenotypic cell profiles is well recognized in contributing to clinical presentations and progression of leishmaniasis. We focused on cutaneous leishmaniasis caused by *L. donovani* that is endemic in our setting, to study the host cytokine and cellular responses across different time points. While being an atypical host-parasite combination relative to global epidemiological patterns, the remarkable uniformity in presentation in the local setting presented a unique opportunity to study the immune responses which are likely key contributors in determining a cutaneous phenotype.

In this study, patient derived cells showed higher levels of secretion of all the inflammatory markers studied (TNF-α, IL-10, TGF-β, IL-12p70, NO) when stimulated with antigen derived from local parasite isolates. The fluctuations in cytokine levels appeared to dampen and stabilize about three days post infection with distinct inflammatory patterns in patients compared to controls. This was further supported by the ROC analysis of the inflammatory markers, with macrophage derived TNF-α, IL-10 and TGF-β showing a clear differentiation between patients and controls at this time point.

We investigated several pro inflammatory markers in this study; TNF-α, NO and IL-12p70. TNF-αplays a crucial role in eliminating intra macrophage parasites through sustained induction of nitric oxide synthase and also promotes development of a protective Th1/IFN-γ response [25]. While NO has been demonstrated to facilitate parasite killing in CL caused by different species of *Leishmania* [26, 27] both TNF-αand NO have been implicated in excessive inflammatory reactions which result in features such as ulceration [28]. IL-12 acts as a vital cytokine which bridges the innate and adaptive immune arms and promotes development of a protective Th1 response [29]. The patient derived cells in our study consistently produced higher levels of NO while TNF-α and IL-12p70 levels were higher at 48 and 72hrs. In a similar study, PBMC derived macrophages from patients with CL and MCL due to *L. braziliensis* demonstrated higher values of TNF-α at 48 hours after infection [23], while another focusing on the same parasite species demonstrated a predominance of TNF-α during active disease [30]. Nitric oxide levels have also shown to be higher in CL by some investigators [31] while others reported no changes [23]. Priming of dendritic cells to secrete IL-12 has been observed in vitro with *L. major* which causes localized cutaneous disease [32], while *L. tropica* which causes more persistent cutaneous lesions did not induce similar responses [24].

Selected anti-inflammatory markers, IL-10 and TGF-β by macrophages and IL-10 by dendritic cells were also evaluated in this study. IL-10 is a well-known cytokine to exert suppressive biological effects on many cell types, including inhibition of macrophage mediated Th1 cell activation [33, 34]. TGF-β is an anti-inflammatory cytokine which predisposes to disease progression and severe manifestations as reported in a variety of infective and inflammatory conditions. In our study IL-10 secreted by both macrophages and dendritic cells were higher in patient derived cells at 72hrs while TGF-β was higher at all three time points. Higher expression of IL-10 in CL and MCL is reported in other studies [35]. A study on role of TGF-β in CL caused by *Leishmania amazonensis, L. donovani chagasi* and *L. braziliensis* has shown that TGF-β led to an increase in parasite numbers when added to in vitro culture [36]. Our results of the comparison of early and late lesions as well as progression to ulceration suggests TGF-β as a key modulator in severity of the lesions. Another study has also demonstrated that TGF-β is associated with chronic forms of the disease [37, 38] or long lasting atypical lesions [39]. It is widely accepted that ulcerated lesions produce more anti-inflammatory cytokines which delays the healing [40].

Overall, our findings showed a mixed inflammatory pattern to be characteristic of the pathogenic process which results in localized CL that we observe in the country. These findings suggest that the effects of anti-inflammatory cytokines which would promote progression of the lesions, is balanced by the pro-inflammatory cytokine responses as well as a sufficient oxidative burst, resulting in limited parasite spread and well localized lesions. Mixed cytokine profiles have also been reported by investigators who conducted similar studies on CL caused by other species [41, 42]. A positive correlation between pro and anti-inflammatory mediators as we observed in dendritic cell responses also are in favour of a regulated response which kills the parasite while limiting tissue injury and is in agreement with reports on similar investigations [43-45]. While in endemic settings the diagnosis of CL is often clinical, aided by microscopy, treated lesions with a poor response or atypical presentations may benefit from complimentary tests based on other biomarkers.

Whereas in vitro studies allow easy manipulation of cells of the immune system and thus simultaneous study of a variety of immune markers, this may not mimic all the conditions *in situ*, in cutaneous lesions. However, our study findings were similar to others who also reported differences in cytokines such as TNF-α, IL-12p70 and IL-10 between local patients and controls by study of skin biopsies [18, 19]. The clinical picture is further supported by cytokine ratios which indicated a greater influence of the pro-inflammatory cytokines. It is also apparent from our results on temporal profiles of expression of cell surface receptors that *Leishmania* parasites cause early activation of macrophages, subsequently inducing polarization towards a M1 phenotype. Polarizing of activated macrophages is influenced by the milieu of cytokines in its micro environment, and the predominance of classically activated macrophages which is a result of exposure to Th1 type cytokines, strengthens our observation of cytokine ratios.

The two groups of controls in this study showed similar cytokine dynamics. While the endemic controls were assumed to have been exposed to infected sandfly bites we did not confirm such exposure by a Montenegro skin test (delayed hypersensitivity testing) primarily due to non-availability of a suitable standardized antigen preparation as well as logistical constraints. Thus, absence or low degree of ‘actual’ exposure could be a contributory factor for this similarity unlike the intermediate profiles reported by others who investigated asymptomatic infections [46, 47]. Another limitation of this study was the relatively small sample size and especially the characteristics of the immune profile of the subgroups with chronic or ulcerated lesions, should be verified in a larger population. The ROC analysis was not extended to differentiating these more severe clinical presentations due to the same reason.

In contrast to a mixed picture as we observed in LCL in the present study, in post-kala-azar dermal leishmaniasis (PKDL), a cutaneous sequel to *L. donovani* infections which cause visceral leishmaniasis, predominance of anti-inflammatory cytokines with M2 polarization of macrophages has been reported [48]. The fact that the infecting parasite species/strain may modulate the immune response in leishmaniasis has been suggested by many experimental and human studies [24, 49]. It is likely that the local parasite also has such antigenic differences when compared to the typical isolates of the species, which has resulted in altered pathogenicity. While previous analyses of genomic sequences by us [50] and others [51] also suggest such differences, a comparison of immune profiles of local patients with LCL and VL would provide more confirmatory evidence on the differential immune responses which influence the phenotype.

In the physiological status, the effects of a cytokine would depend not only on secretory levels but on an interacting network with other cytokines as well as a spectrum of other inflammatory mediators. Effects of other key cytokines in the innate response such as IFN-γ as well as activity of T cells, including cytokine responses of Th17 and T regulatory cells in addition to conventional Th1 and Th2 cells, would need to be explored to obtain a better understanding of the inflammatory dynamics concerned. The early interactions at the host-parasite interface and changes in mediating molecules such as toll-like-receptors (TLRs) [52, 53] are some other factors to be considered in discerning the pathogenic mechanisms.

## Conclusions

Collectively, our data shows that a time dependent mixed inflammatory pattern but with a high pro-inflammatory to regulatory cytokine ratio with classically activated macrophages underlies the clinical phenotype of LCL caused by a dermotropic variant of *L. donovani*. Similar observations of mixed patterns in LCL being reported from different geographical foci suggest that some of the pathogenic mechanisms underlying LCL are conserved across species, thus making these suitable candidates for immune modulatory or immune prophylactic interventions. Our data provides groundwork to evaluate suitability of developing assays based on such serological biomarkers in assessing disease status and prognosis. Further, the study adds to the information on how this intra cellular parasite modulates first line host defenses and provides evidence for likely host factors which prevent visceralization of the parasite.

## Supporting information

**S1 Fig. Comparison of M1/M2 polarization of macrophages derived from patients with acute and chronic lesions**.

A) MI and B) M2) polarization of macrophages stimulated with SLA was compared between patients with acute (duration less than six months) and chronic lesions. The box represents the IQR (i.e. the middle 50% of the observations) with the horizontal line representing the median. The whiskers represent the main body of the data, indicating the range of the data. For statistical analysis, nonparametric Kruskal-Wallis test followed by the Dunn’s multiple comparison test was used (*p<0.05; **p<0.01; ***p<0.001).

**S2 Fig. Comparison of distribution of activated macrophages derived from patients with ulcerated and non ulcerated lesions**.

The distribution of activated macrophages following stimulation with SLA was compared between patients with ulcerated and non ulcerated lesions. The box represents the IQR (i.e. the middle 50% of the observations) with the horizontal line representing the median. The whiskers represent the main body of the data, indicating the range of the data. For statistical analysis, nonparametric Kruskal-Wallis test followed by the Dunn’s multiple comparison test was used (*p<0.05; **p<0.01; ***p<0.001).

## Acknowledgments

The authors wish to thank Dr. KKVN Somarathna, Dr. L. Pathiraja, staff members of the Dermatology Clinics of General Hospitals Anuradhapura, Hambanthota and Padaviya and Dr. Nuwani Manamperi for assistance with patient recruitment, Prof. Neelika Malawige and the staff members of the Department of Immunology and Molecular Medicine, Faculty of Medical Sciences, University of Sri Jayawardenapura for guidance and facilitation of flowcytometry studies and Mr. Sudath Weerasingha for technical assistance.

## Reference

1. World Health Organisation. Leishmaniasis. Available from: https://www.who.int/health-topics/leishmaniasis#tab=tab_1

2. Rosales-Chilama M, Gongora RE, Valderrama L, Jojoa J, Alexander N, Rubiano LC, et al. Parasitological Confirmation and Analysis of Leishmania Diversity in Asymptomatic and Subclinical Infection following Resolution of Cutaneous Leishmaniasis. PLoS Neglected Tropical Diseases. 2015;9(12):e0004273. Available from: https://journals.plos.org/plosntds/article?id=10.1371/journal.pntd.0004273

3. Scharton-Kersten T, Scott P. The role of the innate immune response in Th1 cell development following Leishmania major infection. Journal of Leukocyte Biology. 1995;57(4):515–22. Available from: https://pubmed.ncbi.nlm.nih.gov/7722408/

4. Handman E, Bullen DVR. Interaction of Leishmania with the host macrophage. Trends in Parasitology. 2002;18(8):332–4. Available from: http://www.ncbi.nlm.nih.gov/pubmed/12377273

5. De Almeida MC, Cardoso SA, Barral-Netto M. Leishmania (Leishmania) chagasi infection alters the expression of cell adhesion and costimulatory molecules on human monocyte and macrophage. International Journal of Parasitology. 2003;33(2):153–62. Available from: http://www.ncbi.nlm.nih.gov/pubmed/12633653

6. Donaghy L, Hong HK, Lee HJ, Jun JC, Park YJ, Choi KS. Hemocyte parameters of the Pacific oyster Crassostrea gigas a year after the Hebei Spirit oil spill off the west coast of Korea. Helgoland Marine Research. 2010;64(4):349–55.

7. Rajapaksa US, Ihalamulla RL, Udagedera C, Karunaweera ND. Cutaneous leishmaniasis in southern Sri Lanka. Transactions of Royal Society of Tropical Medicie and Hygiene. 2007;101(8):799–803. Available from: https://academic.oup.com/trstmh/article-lookup/doi/10.1016/j.trstmh.2006.05.013

8. Abeygunasekara PH, Costa YJ, Seneviratne N, Ratnatunga N, Wijesundera MS. Locally acquired visceral leishmaniasis in Sri Lanka. Ceylon Medical Journal. 2007;52(1):30–1. Available from: https://pubmed.ncbi.nlm.nih.gov/17585579/

9. Rathnayake D, Ranawake RR, Sirimanna G, Siriwardhane Y KN and De Silva R. Co-infection of mucosal leishmaniasis and extra pulmonary tuberculosis in a patient with inherent immune deficiency. International journal of dermatology. 2010; 49(5): 549–551

10. Karunaweera ND, Pratlong F, Siriwardane HVYD, Ihalamulla RL, Dedet JP. Sri Lankan cutaneous leishmaniasis is caused by Leishmania donovani zymodeme MON-37. Transactions of Royal Society of Tropical Medicine and Hygiene. 2003;97(4):380–1. Available from: https://pubmed.ncbi.nlm.nih.gov/15259461/

11. Ranasinghe S, Zhang WW, Wickremasinghe R, Abeygunasekera P, Chandrasekharan V, Athauda S, et al. Leishmania donovani zymodeme MON-37 isolated from an autochthonous visceral leishmaniasis patient in Sri Lanka. Pathogens and Global Health. 2012;106(7):421–4. Available from: /pmc/articles/PMC4001626/?report=abstract

12. Mebrahtu YB, Roberts C, Mebrahtu YB, Roberts C, Eys G Van, Guizani I, et al. Human cutaneous leishmaniasis caused by leishmania donovani s.l. in kenya. Transactions of Royal Society of Tropical Medicine and Hygiene. 1993;87(5):598–601.

13. Pratlong F, Bastien P, Perello R, Lami P, Dedet JP. Human cutaneous leishmaniasis caused by Leishmania donovani sensu stricto in Yemen. Transactions of Royal Society of Tropical Medicine and Hygiene. 1995;89(4):398–9. Available from: https://pubmed.ncbi.nlm.nih.gov/7570877/

14. Raina S, Raina RK, Sharma R, Rana BS, Bodh A, Sharma M. Expansion of visceral leishmaniasis to northwest sub-Himalayan region of India: A case series. Journal of Vector Borne Diseases. 2016;53(2):188–91. Available from: http://www.ncbi.nlm.nih.gov/pubmed/27353591

15. Kariyawasam KKGDUL, Selvapandiyan A, Siriwardana HVYD, Dube A, Karunanayake P, Senanayake SASC, et al. Dermotropic Leishmania donovani in Sri Lanka: Visceralizing potential in clinical and preclinical studies. Parasitology. 2018;145(4):443–52. Available from: https://pubmed.ncbi.nlm.nih.gov/29113609/

16. Samaranayake TN, Dissanayake VHW, Fernando SD. Clinical manifestations of cutaneous leishmaniasis in Sri Lanka - Possible evidence for genetic susceptibility among the Sinhalese. Annals of Tropical Medicine and Parasitology. 2008;102(5):383–90. Available from: https://www.tandfonline.com/doi/abs/10.1179/136485908X300779

17. Samaranayake TN, Fernando SD, Dissanayake VHW. Candidate gene study of susceptibility to cutaneous leishmaniasis in Sri Lanka. Tropical Medicine and International Health. 2010;15(5):632–8. Available from: https://pubmed.ncbi.nlm.nih.gov/20214763/

18. Manamperi NH, Oghumu S, Pathirana N, de Silva MVC, Abeyewickreme W, Satoskar AR, et al. In situ immunopathological changes in cutaneous leishmaniasis due to Leishmania donovani. Parasite Immunology. 2017;39(3):e12413. Available from: http://www.ncbi.nlm.nih.gov/pubmed/28112425

19. Galgamuwa LS, Sumanasena B, Iddawela D, Wickramasinghe S, Yatawara L. Assessment of intralesional cytokine profile of cutaneous leishmaniasis caused by Leishmania donovani in Sri Lanka. BMC Microbiology. 2019;19(1):14. Available from: https://bmcmicrobiol.biomedcentral.com/articles/10.1186/s12866-018-1384-4

20. World Medical Association(WMA). WMA Declaration of Helsinki – Ethical Principles for Medical Research Involving Human Subjects. Available from: https://www.wma.net/policies-post/wma-declaration-of-helsinki-ethical-principles-for-medical-research-involving-human-subjects/

21. Barbosa-de-Deus R, Dos Mares-Guia ML, Zacarias Nunes A, Morais Costa K, Gonçalves Junqueira R, Mayrink W, et al. Leishmania major-like antigen for specific and sensitive serodiagnosis of human and canine visceral leishmaniasis. Clinical and Diagnostic Laboratory Immunology. 2002;9(6):1361–6. Available from: https://www.ncbi.nlm.nih.gov/pmc/articles/PMC130090/

22. Lowry OH, Rosebrough NJ, Farr AL, Randall RJ. Protein measurement with the Folin phenol reagent. Journal of Biological Chemistry. 1951;193(1):265–75. Available from: https://pubmed.ncbi.nlm.nih.gov/14907713/

23. Giudice A, Vendrame C, Bezerra C, Carvalho LP, Delavechia T, Carvalho EM, et al. Macrophages participate in host protection and the disease pathology associated with Leishmania braziliensis infection. BMC Infectious Diseases. 2012;12:75. Available from: http://www.ncbi.nlm.nih.gov/pubmed/22458474

24. McDowell MA, Marovich M, Lira R, Braun M, Sacks D. Leishmania priming of human dendritic cells for CD40 ligand-induced interleukin-12p70 secretion is strain and species dependent. Infection and Immunity. 2002;70(8):3994–4001. Available from: https://pubmed.ncbi.nlm.nih.gov/12117904/

25. Wijesuriya H. Correlation between the different cytokines at different time intervals. Figshare [cited 2022 June 2]. Available from: https://figshare.com/s/6d7203ba834cea22639c

26. Wijesuriya H. M2 Type cell polarisation of the macrophages stimulated with the L.donovani. Figshare [cited 2022 June 2]. Available from: https://figshare.com/s/dbd757843c59ae7811bc

27. Wijesuriya H. Maturation of macrophage cells stimulated with Leishmania donovani antigen. Figshare [cited 2022 June 2]. Available from: https://figshare.com/s/04b6424f668c0Ca4cafe

28. Wijesuriya H. Production of IL-10 and IL12p70 by macrophages stimulated with L.donovani antigen with the severity. Figshare [cited 2022 June 2]. Available from: https://figshare.com/s/878c43b485d2fe8a76ee

29. Umakant Sharma SS. Immunobiology of leishmaniasis. Indian Journal of Experimental Biology. 2009; 47(6): 412–423 Available from: https://pubmed.ncbi.nlm.nih.gov/19634705/

30. Serarslan G, Atik E. Expression of inducible nitric oxide synthase in human cutaneous leishmaniasis. Molecular and Cellular Biochemistry. 2005;280(1–2):147–9. Available from: https://pubmed.ncbi.nlm.nih.gov/16311916/

31. Kumar R, Bumb RA, Salotra P. Evaluation of localized and systemic immune responses in cutaneous leishmaniasis caused by Leishmania tropica: Interleukin-8, monocyte chemotactic protein-1 and nitric oxide are major regulatory factors. Immunology. 2010;130(2):193–201. Available from: https://pubmed.ncbi.nlm.nih.gov/20102417/

32. Carneiro PP, Conceição J, Macedo M, Magalhães V, Carvalho EM, Bacellar O. The role of nitric oxide and reactive oxygen species in the killing of Leishmania braziliensis by monocytes from patients with cutaneous leishmaniasis. PLoS One. 2016;11(2):e0148084 Available from: pmc/articles/PMC4739692/?report=abstract

33. Martínez-López M, Iborra S, Conde-Garrosa R, Sancho D. Batf3-dependent CD103+ dendritic cells are major producers of IL-12 that drive local Th1 immunity against Leishmania major infection in mice. European Journal of Immunology. 2015;45(1):119–29. Available from: https://pubmed.ncbi.nlm.nih.gov/25312824/

34. Souza MA, Castro Mcab, Oliveira AP, Almeida AF, Reis LC, Silva CJ, et al. American Tegumentary Leishmaniasis: Cytokines and Nitric Oxide in Active Disease and After Clinical Cure, With or Without Chemotherapy. Scandinavian Journal of Immunology. 2012;76(2):175–80. Available from: http://www.ncbi.nlm.nih.gov/pubmed/22537157

35. Dayakar A, Chandrasekaran S, Kuchipudi S V., Kalangi SK. Cytokines: Key determinants of resistance or disease progression in visceral leishmaniasis: Opportunities for novel diagnostics and immunotherapy. Frontiers in Immunology. 2019;10:670. Available from: www.frontiersin.org

36. Bacchetta R, Gambineri E, Roncarolo MG. Role of regulatory T cells and FOXP3 in human diseases. Journal of Allergy and Clinical Immunology. 2007;120(2);227–235. Available from: http://www.jacionline.org/article/S0091674907012432/fulltext

37. Pirmez C, Yamamura M, Uyemura K, Paes-Oliveira M, Conceicao-Silva F, Modlin RL. Cytokine patterns in the pathogenesis of human Leishmaniasis. Journal of Clinical Investigation. 1993;91(4):1390–5. Available from: /pmc/articles/PMC288111/?report=abstract

38. Melby PC, Andrade-Narvaez FJ, Darnell BJ, Valencia-Pacheco G, Tryon V V., Palomo-Cetina A. Increased expression of proinflammatory cytokines in chronic lesions of human cutaneous leishmaniasis. Infection and Immunity. 1994;62(3):837–42. Available from: /pmc/articles/PMC186190/?report=abstract

39. Hejazi SH, Hoseini SG, Javanmard SH, Zarkesh SH, Khamesipour A. Interleukin-10 and transforming growth factor-β in early and late lesions of patients with Leishmania major induced cutaneous Leishmaniasis. Iranian Journal of Parasitology. 2012;7(3):16–23. Available from: /pmc/articles/PMC3469188/?report=abstract

40. Diaz NL, Zerpa O, Ponce LV, Convit J, Rondon AJ, Tapia FJ. Intermediate or chronic cutaneous leishmaniasis: leukocyte immunophenotypes and cytokine characterisation of the lesion. Experimental Dermatology. 2002;11(1):34–41. Available from: http://doi.wiley.com/10.1034/j.1600-0625.2002.110104.x

41. Castellano LR, Filho DC, Argiro L, Dessein H, Prata A, Dessein A, et al. Th1/Th2 immune responses are associated with active cutaneous leishmaniasis and clinical cure is associated with strong interferon-γ production. Human Immunology. 2009;70(6):383–90.

42. Baratta-Masini A, Teixeira-Carvalho A, Malaquias LCC, Mayrink W, Martins-Filho OA, Corrêa-Oliveira R. Mixed cytokine profile during active cutaneous leishmaniasis and in natural resistance. Frontiers in Bioscience; 2007;12:839–49. Available from: https://pubmed.ncbi.nlm.nih.gov/17127341/

43. Antonelli LRV, Dutra WO, Almeida RP, Bacellar O, Gollob KJ. Antigen specific correlations of cellular immune responses in human leishmaniasis suggests mechanisms for immunoregulation. Clinical Experimental Immunology. 2004;136(2):341–8. Available from: /pmc/articles/PMC1809031/?report=abstract

44. Oliveira F, Bafica A, Rosato AB, Favali CBF, Costa JM, Cafe V, et al. Lesion size correlates with Leishmania antigen-stimulated TNF-levels in human cutaneous leishmaniasis. American Journal of Tropical Meicine and Hygiene. 2011;85(1):70–3. Available from: /pmc/articles/PMC3122347/?report=abstract

45. Follador I, Araújo C, Bacellar O, Araújo CB, Carvalho LP, Almeida RP, et al. Epidemiologic and immunologic findings for the subclinical form of Leishmania braziliensis infection. Clinical Infectious Diseases. 2002;34(11). Available from: https://pubmed.ncbi.nlm.nih.gov/12015707/

46. Ramos PK, Carvalho KI, Rosa DS, Rodrigues AP, Lima LV, Campos MB, et al. Serum cytokine responses over the entire clinical-immunological spectrum of human Leishmania (L.) infantum chagasi infection. Biomed Research International. 2016;2016. 6937980. Available from: https://www.ncbi.nlm.nih.gov/pmc/articles/PMC4802012/

47. Mukhopadhyay D, Mukherjee S, Roy S, Dalton JE, Kundu S, Sarkar A, et al. M2 Polarization of Monocytes-Macrophages Is a Hallmark of Indian Post Kala-Azar Dermal Leishmaniasis. PLoS Neglected Tropical Diseases. 2015;9(10):2. Available from: /pmc/articles/PMC4619837/?report=abstract

48. Reed SG. TGF-beta in infections and infectious diseases. Microbes and Infection. 1999;1(15):1313–25. Available from: http://www.ncbi.nlm.nih.gov/pubmed/10611760

49. Samarasinghe SR, Samaranayake N, Kariyawasam UL, Siriwardana YD, Imamura H, Karunaweera ND. Genomic insights into virulence mechanisms of Leishmania donovani: Evidence from an atypical strain. BMC Genomics. 2018;19(1):843. Available from: https://bmcgenomics.biomedcentral.com/articles/10.1186/s12864-018-5271-z

50. Zhang WW, Ramasamy G, McCall L-I, Haydock A, Ranasinghe S, Abeygunasekara P, et al. Genetic Analysis of Leishmania donovani Tropism Using a Naturally Attenuated Cutaneous Strain. PLoS Pathogens. 2014;10(7):e1004244. Available from: https://dx.plos.org/10.1371/journal.ppat.1004244

51. Tolouei S, Hejazi SH, Ghaedi K, Khamesipour A, Hasheminia SJ. TLR2 and TLR4 in Cutaneous Leishmaniasis Caused by Leishmania major. Scandinavian Journal of Immunology. 2013;78(5):478–84. Available from: http://doi.wiley.com/10.1111/sji.12105

52. Ajdary S, Ghamilouie MM, Alimohammadian MH, Riazi-Rad F, Pakzad SR. Toll-like receptor 4 polymorphisms predispose to cutaneous leishmaniasis. Microbes and Infecttion. 2011;13(3):226–31. Available from:

53. Tolouei S, Hejazi SH, Ghaedi K, Khamesipour A, Hasheminia SJ. TLR2 and TLR4 in Cutaneous Leishmaniasis Caused by Leishmania major. Scandinavian Journal of Immunology. 2013;78:478–84. doi: 10.1111/sji.12105.

